# Performance of genetic imputation across commercial crop species

**DOI:** 10.1101/2021.12.01.470712

**Authors:** Steve Thorn, Andrew Whalen, Sonja Kollers, Mahmood Gholami, Helena Sofia da Silva, Valentin Wimmer, John M Hickey

**Author notes:** Email addresses: ST AW SK MG HSdS VW JMH.

## Abstract

We show that accurate imputation can be carried out in three commercial plant species (maize, sugar beet and wheat) and that accurate imputation does not require a pedigree, although pedigree information can improve accuracy and speed. Our approach uses a hidden Markov model to build a haplotype library from individuals genotyped at high-density and then uses this library to impute low-density genotyped individuals to high-density. To build the library, we use founders when the pedigree is known, or a sample of progeny when the pedigree is unknown. Without a pedigree, and with 50 individuals genotyped at high-density and 100 low-density markers per chromosome, the median accuracies were 0.97 (maize), 0.96 (sugar beet), and 0.94 (wheat). We obtained similar accuracies with a pedigree. For biparental crosses with 100 markers per chromosome, median accuracies were 0.96 (maize), 0.96 (sugar beet) and 0.94 (wheat). For the imputation scenarios without a pedigree, we compared accuracies with those obtained by running Beagle 5.1. In all but one scenario, our method outperformed Beagle. We believe that plant breeders can effectively apply imputation in many crop species.

## Introduction

In this paper, we show that accurate imputation can be carried out in three commercial plant species (maize, sugar beet and wheat) and that accurate imputation does not require a pedigree, although pedigree information can improve accuracy and speed. Imputation is a valuable and widely used tool in plant breeding programmes and can decrease the cost of genomic selection. Genomic selection can increase the effectiveness of plant breeding [1,2] but requires genotyping large numbers of individuals at large-enough marker density, and this can be prohibitively expensive. One strategy to reduce cost is to genotype only a small number of individuals at high-density (higher unit cost) and a much larger number at low-density (lower unit cost). Imputation can then recover much of the lost information by finding haplotypes common to the low- and high-density genotypes and allows genomic selection to be applied [3–5].

A common way to perform imputation is to use a hidden Markov model, which models an individual’s genotype as a mosaic of haplotypes from a haplotype library (built from individuals genotyped at high-density). We examine two ways to build the haplotype library. For accurate imputation, the library should contain as many of the (unique) haplotypes present in the population as possible. A guaranteed way to build such a library is to build it from founder genotypes; however, these are not always available. If the founders are not available, we can use a sample of individuals from the population. Increasing the number of sampled individuals will increase the ‘completeness’ of the haplotype library (and hence increase imputation accuracy), albeit at an increased cost of genotyping. We refer to the two ways to build the haplotype library as imputation *with a pedigree* (founders used to build the library) or *without a pedigree* (sample of progeny used to build the library).

One of the steps to applying imputation for plant breeding programs is the demonstration that already-existing imputation methods can yield good accuracies in commercial plant breeding programs. Previous work by Pook et al. 2019 [6], found that several versions of an already existing software package, Beagle (versions 4.0, 4.1, 5.0 and 5.1 [7–10]), achieved good imputation accuracies but depended on setting model parameters far away from their defaults to take into account the lower genetic diversity present in plant and animal breeding populations.

In this study, we evaluate the performance of a hidden Markov model approach on commercial breeding material across three crop species: maize (*Zea mays* L.), sugar beet (*Beta vulgaris* L.) and wheat (*Triticum aestivum* L.).

We obtained high imputation accuracies even with few low-density markers. Without a pedigree, and with 50 individuals genotyped at high-density and 100 low-density markers per chromosome, the median accuracies were 0.97 (maize), 0.96 (sugar beet), and 0.94 (wheat). We obtained similar accuracies with a pedigree. For the biparental crosses with 100 markers per chromosome, median accuracies were 0.96 (maize), 0.96 (sugar beet) and 0.94 (wheat). Imputation accuracy was better with a pedigree, but only when similar numbers of individuals were used to build the haplotype libraries.

We believe that plant breeders can effectively apply imputation in many crop species and that while pedigree information is not required to obtain good accuracy, it can increase the accuracy and speed of imputation.

## Materials and Methods

### Imputation method

We used a Li and Stephens style hidden Markov model to perform imputation and phasing with and without a pedigree. This imputation method consists of two parts. First, we create a haplotype library by phasing the genotypes of individuals genotyped at high-density. Then we use the haplotype library to impute low-density genotyped individuals to high-density. For imputation with a pedigree, we restricted the haplotype library to founder haplotypes and examined biparental, three and four-way crosses separately.

### Hidden Markov model

To perform imputation, we used a Li and Stephens hidden Markov model [11]. Under this model, we assume that an individual’s haplotypes are a mosaic of haplotypes from a haplotype reference library. The model infers which haplotypes the individual carries at each locus. We can define the model to work either on a single chromosome in a haploid model (for inbred individuals) or on pairs of chromosomes in a diploid model (for outbred individuals). The states in the model are either single haplotypes (haploid model) or pairs of haplotypes (diploid model) from the reference library. To define the hidden Markov model, we construct a state transition function, *P*(*S*_*i*_|*S*_*i*−1_), and an emission function, *P*(*G*_*i*_|*S*_*i*_), where *S*_*i*_ is the hidden state and *G*_*i*_ the genotype at locus *i*. The transition function describes how states change throughout the genome, based on ancestral recombinations. The emission function connects the haplotype states to the observed genotype data, which may be collected with some error. The form of these functions depends on the type of hidden Markov model.

### Haploid hidden Markov model

The haploid hidden Markov model assumes the genotype is homozygous at all loci (we set any residual heterozygous loci to missing). Homozygous loci are appropriate for inbred individuals generated either from multiple rounds of selfing or by doubled haploid technology. It models a target genotype as a mosaic of single haplotypes. Each hidden state corresponds to one of these haplotypes: if there are *H* haplotypes, then there are *H* possible hidden states. Initially, each state is deemed equally likely.

The transmission function is defined by examining the possibility of moving to another state as we move from one locus to the next. The model assumes that if a transition between states occurs, then all destination states are equally likely, *including* a transition back to the original state. We follow Li et al., 2010 [12], and define this state-to-state transition probability as *θ/H*, where *H* is the number of states (haplotypes), and *θ* is a model parameter. The transition function is then:

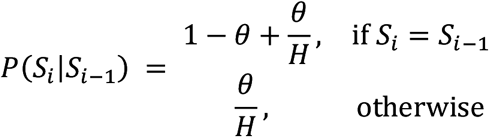

In this model, *θ* represents an ancestral recombination rate, which gives the likelihood of a recombination occurring between the source haplotype in the haplotype reference library and the individual’s haplotype. For imputation with a pedigree, this can be calculated by using the genetic map length. We approximate this by using *θ* = 1*/N*, where *N* is the number of high-density loci in the chromosome being imputed. Such a value corresponds to a single recombination (on average) between a parent and offspring, or equivalently a chromosome length of 100 cM. Different values might be more appropriate for imputation without a pedigree, or for larger or smaller chromosomes. In practice, however, we found that imputation accuracy is not particularly sensitive to this parameter and we used a fixed value of *θ* = 1*/N* for all loci. We do not iteratively refine it, as do Li et al., 2010.

We model the emission probabilities using an error-rate parameter *ϵ*. The emission probability is 1 − *ϵ* if the state’s allele matches that of the target haplotype and *ϵ* if not. If the target allele is missing, the emission probability is set such that all states are equally likely (probability 1*/H*). We also found that imputation accuracy is not sensitive to this parameter and used a fixed value of 0.01 across all loci.

### Diploid hidden Markov model

The diploid hidden Markov model is similar to the haploid one. Instead of focusing on a single haplotype, it represents a target genotype as a mosaic of *pairs* of haplotypes. This model is appropriate for outbred individuals. The hidden states correspond to all possible ordered pairs of haplotypes, such that *H* haplotypes correspond to *H*^2^ states. As with the haploid model, all possible states are initially equally likely.

The probability of changing state from one locus to the next depends on the kind of state change. Assuming that the transitions of each haplotype (in the state) are independent, we can calculate the transition function by simply multiplying those from the haploid model:

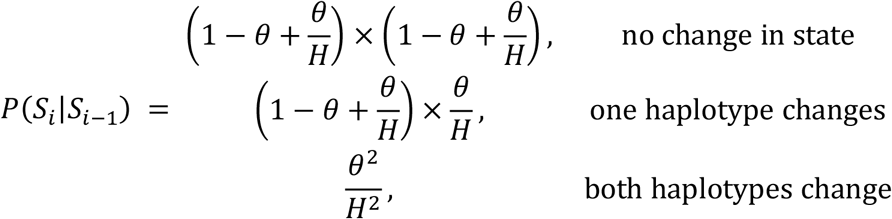

As with the haploid model, we set *θ* to 1*/N* across all loci.

We simplify the emission probabilities for the diploid case compared to Li et al., 2010, and only consider cases where the state’s pair of alleles is consistent with the target genotype (with probability 1 − *ϵ*) or not (probability *ϵ*). If the target genotype is missing, we set the emission probability to 1*/H*^2^ such that all states are equally likely. We used a fixed value of 0.01 for *ϵ*.

### Phasing and imputing genotypes

We calculate the probability of each state using the forward-backward algorithm (see, e.g., [13]), which gives a probability distribution, over the possible states, at each locus. We calculate genotype dosages by converting the state distributions, at each locus, to distributions over possible genotypes (0, 1, or 2). At a given locus, each state corresponds to a genotype, and we calculate the probability for each genotype by summing over the relevant state probabilities. The genotype dosage is the expected value of the probability distribution over genotypes. We calculate dosages in the imputation part of the method.

For a target genotype, we sample states, in a similar way, by applying the forward algorithm, but sampling on the backwards pass (see Thompson [14], pp. 95 for an example of this). Each sample is one of the possible mosaics of haplotypes that models the genotype (single haplotypes in the haploid model or pairs of haplotypes in the diploid model). We use sampling in the haplotype library creation part of the method.

### Haplotype filtering

For the diploid hidden Markov model, it is often not computationally tractable to consider all possible states, as the number of states grows as *H*^2^. To control the complexity, we reduce the number of haplotypes to a (fixed-size) subset of haplotypes that are most similar to the target genotype. (We also keep this as an option for the haploid model, but it is often not necessary as the size of the state space only grows linearly). We divide each chromosome into windows containing equal numbers of markers. In each window, we rank the haplotypes in order of increasing number of opposite homozygous markers compared to the target genotype. We then select the top-ranked haplotypes from these lists. In the event of tied ranks, we randomly choose from the tied haplotypes. This approach is similar to that used in IMPUTE2 [15], except we do not iteratively refine the haplotypes selected based on the previous phasing of the haplotypes.

For all the studies presented here, we used five windows per chromosome to filter a subset of 100 haplotypes from the haplotype library (unless the library contained fewer haplotypes, in which case we used all of them).

### Haplotype library creation

To create a haplotype library, we use a modified version of the method of Li et al., 2010 [12]. We only present a summary here and refer the reader to the reference for full details. First, we select individuals that will make up the haplotype library. Usually, these are individuals genotyped at high-density. The method phases these individuals using an iterative process as follows:

i. randomly phase each genotype into a pair of haplotypes
ii. sample a (refined) pair haplotypes using a suitable hidden Markov model
iii. correct and update the haplotype pair
iv. repeat steps ii) and iii) a fixed number of times.

In pilot simulations, we found that good imputation accuracy required only a few iterations (5–10) and we used five iterations for all the studies here.

#### Random phasing

We randomly phase each target genotype into a pair of haplotypes. At heterozygous loci, the major allele is randomly assigned to either the first or second haplotype with equal probability. Missing loci are filled in according to the population minor allele frequency, MAF_*i*_: at locus *i*, each haplotype is assigned the minor allele with probability MAF_*i*_, or the major allele with probability 1 − MAF_*i*_. Homozygous loci are de-facto phased.

#### Sample and correct haplotypes

We construct a hidden Markov model for each target genotype we are phasing. For a given target, the reference haplotypes we use are a filtered subset of haplotypes from all other individuals in the library (i.e., we exclude the haplotypes from the target individual). We sample haplotypes from the model to generate a pair of haplotypes. In the case of the haploid model, the pair of haplotypes is formed from two identical copies of the single sampled haplotype. A sample from the diploid model is already a pair of haplotypes. We correct the sampled pair of haplotypes to be consistent with the target genotype. At inconsistent loci, we alter the haplotypes’ alleles using the same random phasing strategy as already described.

We repeat the sampling and correction steps for each individual in the library, and then update the library to consist of the new haplotype pairs. Each time we sample (and correct) a pair of haplotypes, we refine the phasing.

### Imputation of low-density genotyped individuals

The method imputes low-density genotyped individuals to high-density using the same hidden Markov model and high-density haplotypes from a haplotype library. The hidden Markov model represents a low-density genotype as a mosaic of high-density haplotypes. We define the model over the high-density markers so that the state probability distributions are also defined over these markers. We pad the low-density genotypes with missing markers such that they have the same number of markers (as the high-density ones). From the probability distributions, we calculate genotype dosages at all high-density marker positions, effectively imputing the individual’s low-density genotype at the high-density loci. If needed, imputed genotypes can be calculated by rounding the dosages to the nearest integer.

### Plant materials

We used data from three crop species: maize (*Zea mays* L.), sugar beet (*Beta vulgaris* L.) and wheat (*Triticum aestivum* L.). Maize data consisted of 338 double haploid (DH) progeny generated from crosses of 26 DH lines, with the crosses being a mix of biparental, three- and four-way. Sugar beet data consisted of 1614 lines from biparental crosses of 41 heterozygous parents. The progeny were selfed twice. Wheat data consisted of 1035 DH lines from crosses of 113 inbred lines, with the crosses being a mix of biparental and three-way. Parents and progeny were genotyped on crop-specific marker arrays with different numbers of markers. This study used approximately 600k markers for maize, 20k for sugar beet and 25k for wheat. Table 1 shows the numbers of genotyped individuals used in the imputation studies.

**Table 1:** numbers of genotyped individuals used in the *with* and *without a pedigree* imputation studies. Note: for sugar beet and wheat, we used more individuals in the without a pedigree case as we could use individuals irrespective of whether their founders were genotyped.

|  | Without a<br>pedigree | With a pedigree |  |  |  |  |  |
| --- | --- | --- | --- | --- | --- | --- | --- |
|  | Biparental<br>Progeny | Biparental |  | Three-way |  | Four-way |  |
|  |  | Founders | Progeny | Founders | Progeny | Founders | Progeny |
| Maize | 127 | 7 | 127 | 10 | 142 | 13 | 69 |
| Sugar beet | 1614 | 32 | 1338 | – | – | – | – |
| Wheat | 555 | 60 | 506 | 71 | 443 | – | – |

All materials described in this study are proprietary to KWS SAAT SE & Co. KGaA.

### Imputation scenarios without a pedigree

To test the accuracy of imputation without a pedigree, we used all biparental cross data from the three crops but excluded the parents. We varied the numbers of individuals genotyped on the high-density array to be 10, 20, 50, 100, 200, 500, or 1000. The remaining individuals were assigned to a simulated, low-density array with 10, 50, 100, 250, 500 or 1000 markers per chromosome. To account for the effect of which individuals are genotyped at high-density, we created ten replicates by repeating the partitioning of individuals into the high-and low-density arrays.

To perform the imputation, we used the individuals genotyped at high-density to build the haplotype library and imputed the low-density genotypes to high-density. We compared both haploid and diploid hidden Markov models with the expectation that the haploid model would be better suited to maize and wheat and the diploid model to sugar beet. In practice, we found the diploid model generally outperformed the haploid one even for maize and wheat, and so report results only for the diploid model.

### Imputation scenarios with a pedigree

When the pedigree is known, we imputed progeny genotyped at low-density from their parents genotyped at high-density. We used data from biparental crosses (maize, sugar beet and wheat), three-way crosses (maize and wheat) and four-way crosses (maize only). As with the scenarios without a pedigree, we simulated six low-density marker arrays for the progeny with 10, 50, 100, 250, 500 or 1000 markers per chromosome.

We used all high-density, parent genotypes to build the haplotype library and imputed the progeny by restricting the haplotype reference library to just the individual’s parent haplotypes. We also compared the haploid and diploid hidden Markov models and found that the diploid model generally outperformed the haploid one, and so report results only for the diploid model.

### Simulated marker arrays

We simulated low-density marker arrays by choosing a fixed number of markers from the original data. We followed the method described in Whalen et al., 2019 [16] and selected markers that have high minor allele frequency and are approximately evenly-spaced.

To simulate a low-density array with *n* markers per chromosome, we divide each high-density chromosome into *n* evenly-spaced windows and score each marker with the following function:

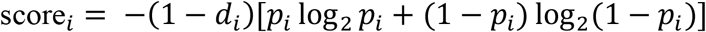

where *p*_*i*_ is the minor allele frequency of marker *i* and *d*_*i*_ is the fractional distance between the marker and the centre of its window. This function is maximised for markers that are close to the centre of their window (1 − *d*_*i*_ ≈ 1) and have minor allele frequency close to 0.5. For each window, we chose the marker with the highest score to be included on the low-density array.

### Imputation accuracy

For each scenario, we imputed the individuals one chromosome at a time. Imputation accuracy was measured using the Pearson correlation between the true genotype and imputed genotype dosages both corrected for minor allele frequency (see, e.g., [17,18]). To obtain an accuracy for each chromosome, we use the median accuracy over all imputed individuals and all ten replicates. For wheat, we did not attempt to impute the D genome due to the low numbers of assigned markers (average of 54.7 markers per chromosome).

### Comparison with Beagle

For the biparental cross scenarios without a pedigree, we compared accuracies with those obtained by running Beagle 5.1 [7,10]. We ran Beagle in two steps: first creating a haplotype library from the individuals genotyped at high-density and then imputing the individuals genotyped at low-density using that haplotype library. In both steps, we used the default settings, except we used an effective population size of 100, following Pook et al. [6].

## Results

Increasing the number of low-density markers increases imputation accuracy. For imputation without a pedigree, increasing the number of individuals used to build the haplotype library increases accuracy. Imputation with a pedigree is more accurate than without a pedigree, when the same number of individuals (founders or progeny) are used to build the haplotype libraries. The accuracy of our method is better than Beagle in the majority of cases.

### Imputation accuracy without a pedigree

Figure 1 shows imputation accuracy for biparental cross scenarios without a pedigree, versus the number of markers on the low-density array when 50 individuals were used to build the haplotype library (100 haplotypes). This figure, and subsequent box plots, show the variation in accuracy across chromosomes: the data for each box are the per chromosome median accuracies. Accuracies increase smoothly with increasing marker density and plateau at around 250 markers. The accuracies increase from 0.85 (maize), 0.77 (sugar beet) and 0.69 (wheat) at ten markers to 0.99 (maize), 0.98 (sugar beet) and 0.97 (wheat) at 250 markers.

**Figure 1:**
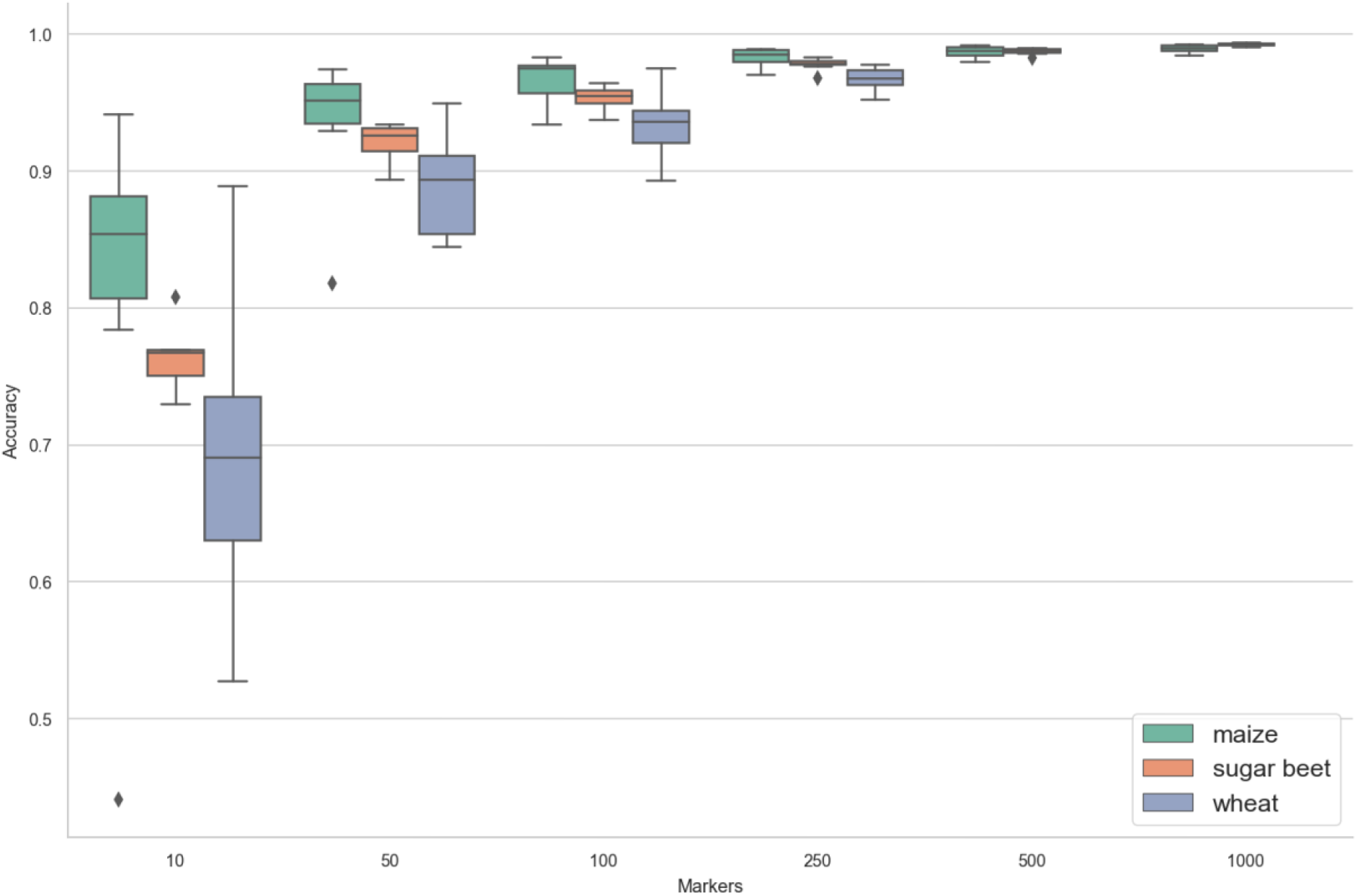
imputation accuracy versus number of low-density markers (per chromosome) for biparental crosses of maize, sugar beet and wheat. The reference haplotype library contains 100 haplotypes (built from 50 individuals genotyped at high-density). Note: there are no boxes for wheat at 500 or 1000 markers as all the wheat chromosomes have fewer than 500 high-density markers. The data for each box are the per chromosome median accuracies. Model parameters, as described in the text.

For imputation without a pedigree, the number of individuals used to build the haplotype library is an important factor. Increasing the number of individuals increases imputation accuracy, but also increases genotyping cost. Figure 2 shows imputation accuracy versus the number of individuals used to build the library for 100 low-density markers. Accuracies increase smoothly as the number of individuals increases, reaching a plateau after approximately 100 individuals.

**Figure 2:**
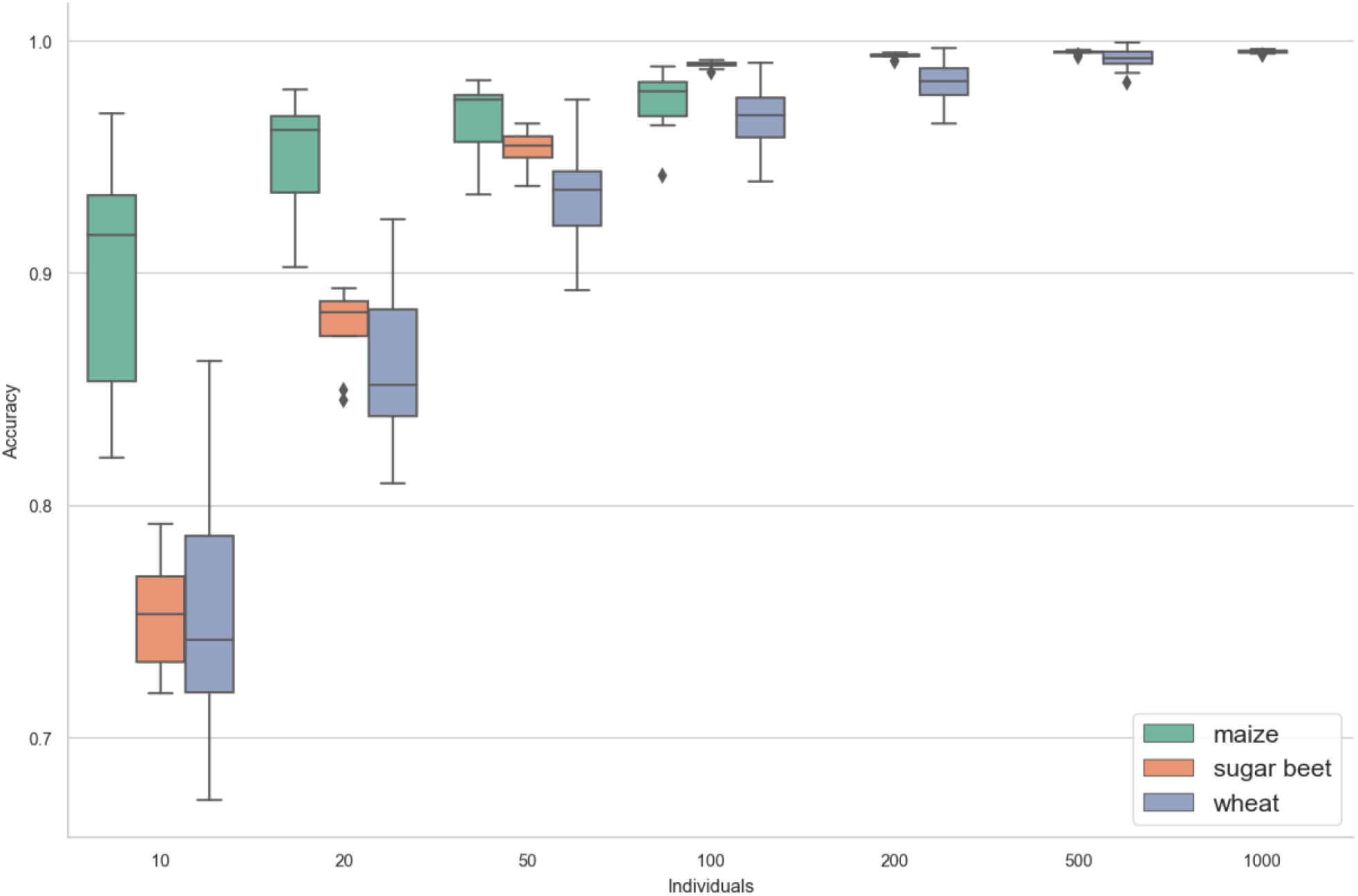
imputation accuracy versus the number of individuals used to build the haplotype library (genotyped at high-density) for biparental crosses of maize, sugar beet and wheat. The number of low-density markers is fixed at 100. Note: there are no boxes for maize (200, 500 or 1000 individuals) or wheat (1000 individuals) as the corresponding datasets contain fewer individuals. The data for each box are the per chromosome median accuracies. Model parameters, as described in the text.

### Imputation accuracy with a pedigree

Results for imputation with a pedigree are similar. Figure 3 shows imputation accuracy versus the number of markers on the low-density array for maize (biparental, three-way and four-way crosses), sugar beet (biparental only) and wheat (biparental and three-way). Accuracies increase smoothly and reach a plateau at around 250 markers. In all cases, imputation accuracy decreases with increasing numbers of founders. For biparental crosses, the accuracies increase from 0.80 (maize) 0.79 (sugar beet), 0.72 (wheat) at ten markers to 0.97 (maize), 0.98 (sugar beet), 0.97 (wheat) at 250 markers. For three-way crosses, accuracies increase from 0.72 (maize), and 0.66 (wheat) at ten markers to 0.96 (maize) and 0.96 (wheat) at 250 markers. Accuracies for the four-way cross (maize only) increase from 0.48 at ten markers to 0.91 at 250 markers.

**Figure 3:**
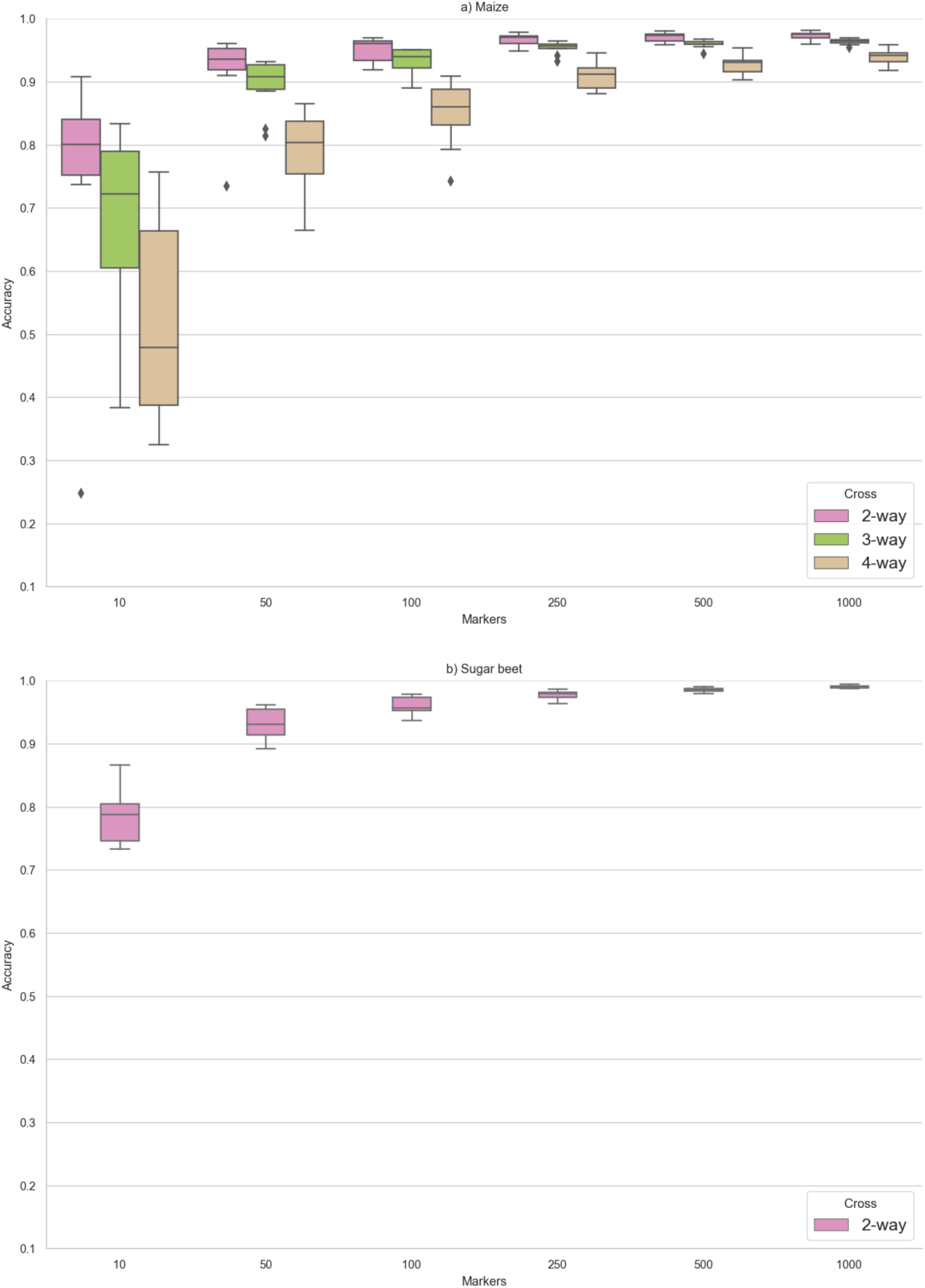
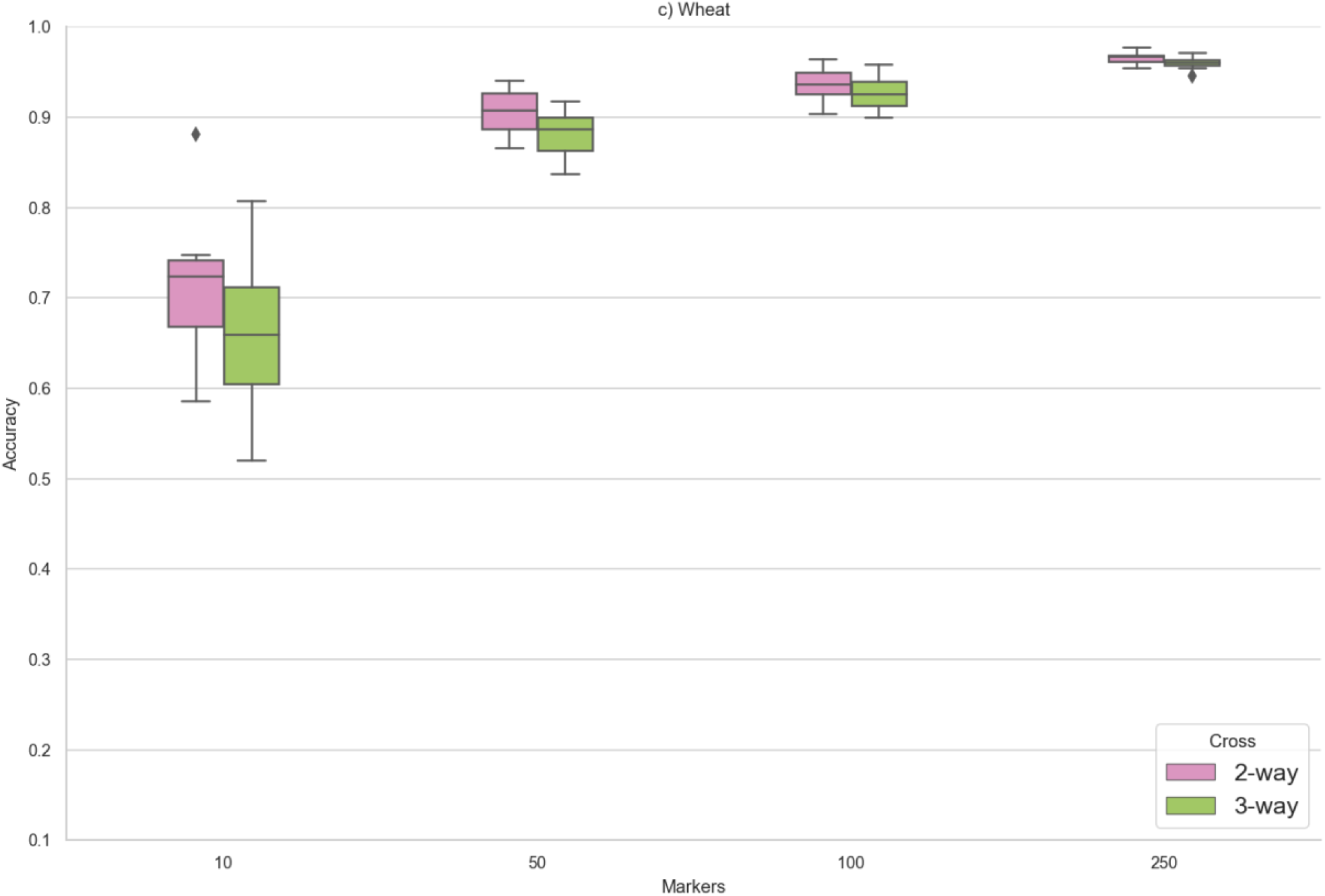
imputation accuracy versus number of low-density markers (per chromosome) for a) maize (biparental, three- and four-way crosses), b) sugar beet (biparental) and c) wheat (biparental and three-way). Note: there are no boxes for wheat at 500 or 1000 markers as all wheat chromosomes have fewer than 500 high-density markers. The data for each box are the per chromosome median accuracies. Model parameters, as described in the text.

### Comparison between imputation with and without a pedigree

Imputation accuracy with a pedigree is higher when a similar number of individuals are used to build the haplotype library. Figure 4 compares the imputation accuracy between scenarios for biparental crosses with and without a pedigree. For all crops, the accuracy of the scenarios with a pedigree is greater when the number of individuals used to build the haplotype libraries is comparable.

**Figure 4:**
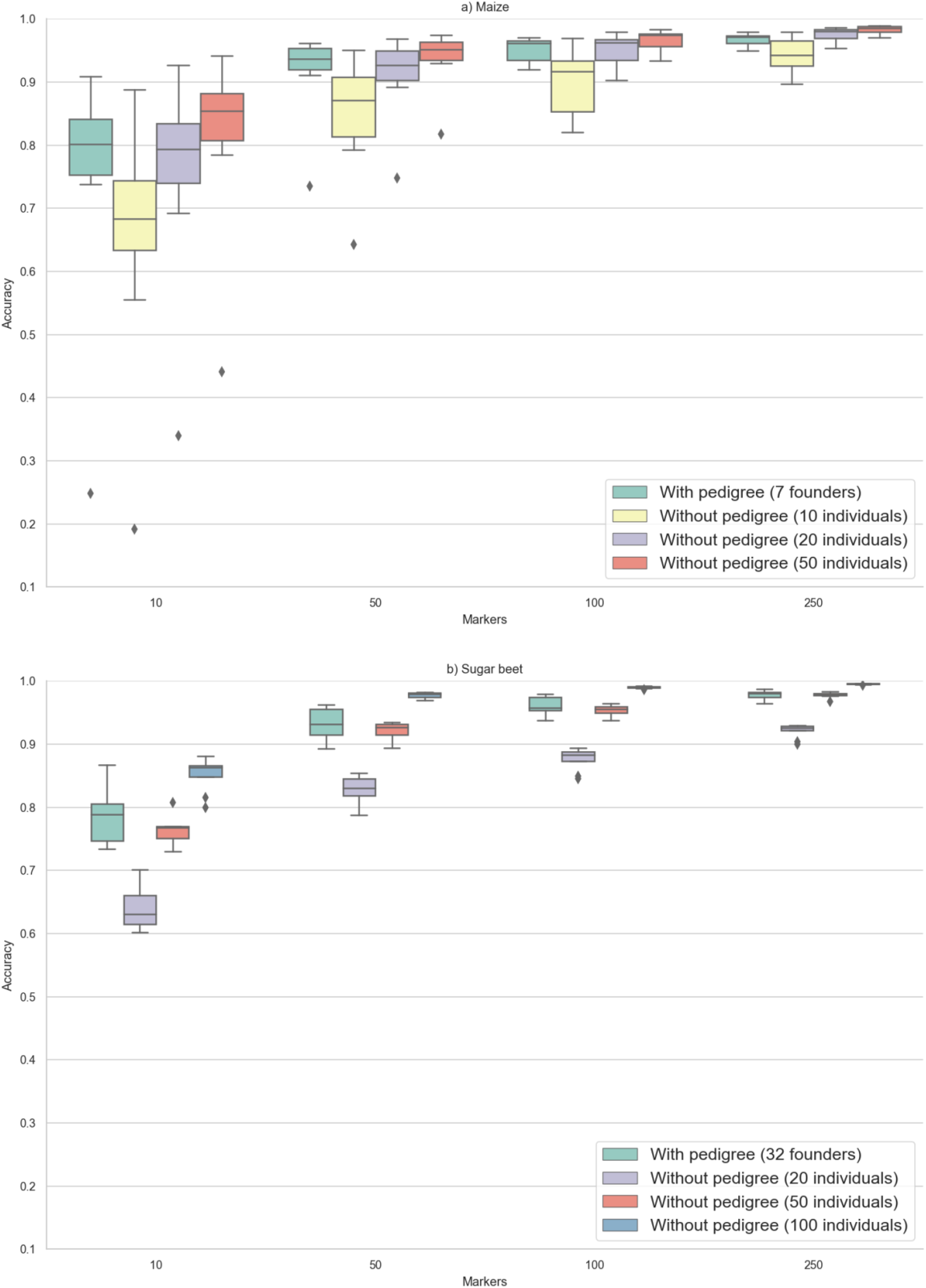
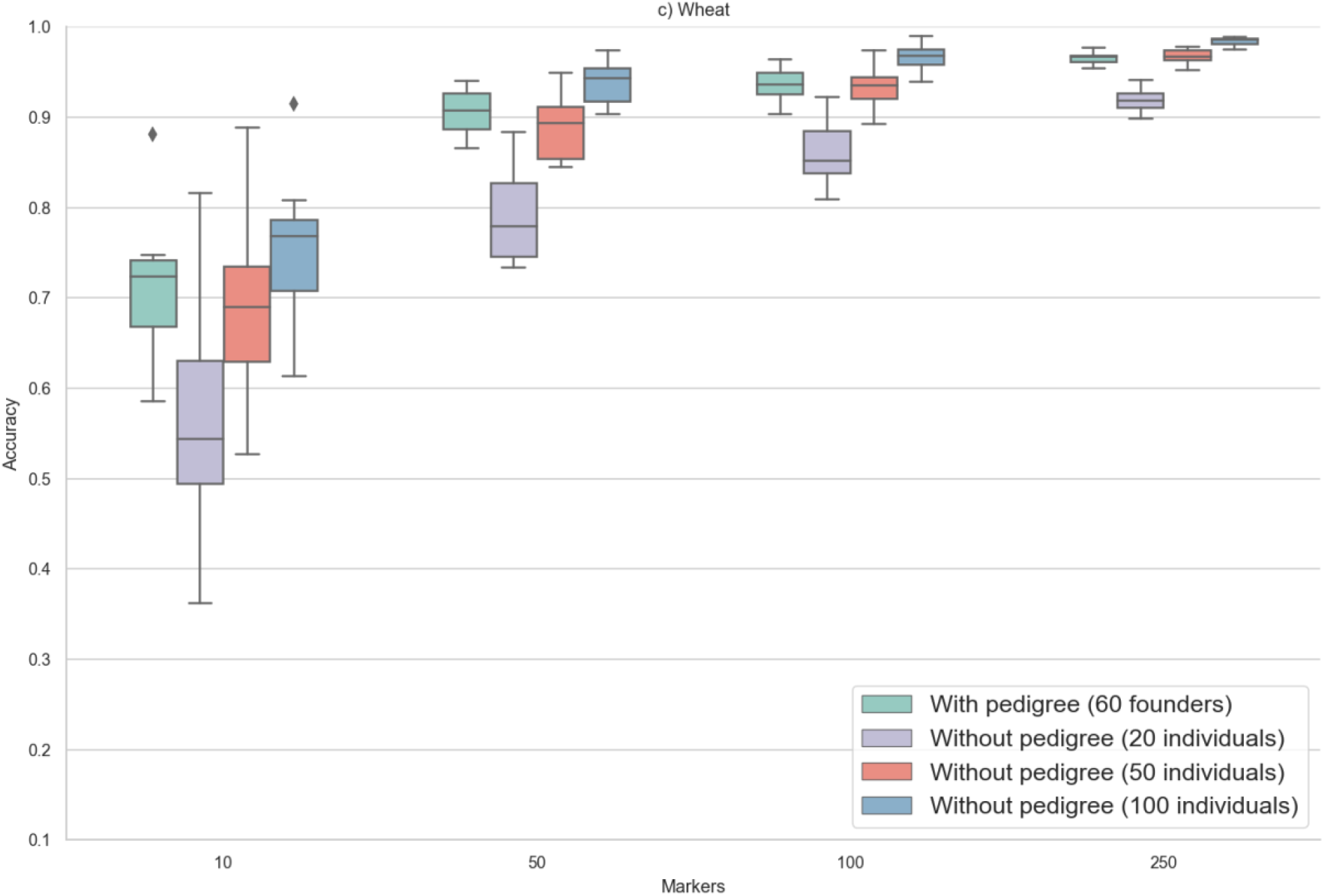
imputation accuracy versus number of low-density markers (per chromosome) with a pedigree (all founders used to build the haplotype library) and without a pedigree (10, 20, 50 or 100 individuals used to build the haplotype library). The data for each box are the per chromosome median accuracies. Model parameters, as described in the text.

It is possible to exceed the accuracy of imputation with a pedigree by using enough individuals (compared to founders) to build the haplotype library. In maize, imputation using 50 progeny to build the haplotype library has higher accuracy than imputation with a pedigree, in which seven founders are genotyped at high density. For the other two crops, the corresponding numbers, at which the imputation accuracy is exceeded, are 100 progeny versus 32 founders for sugar beet and 100 progeny versus 60 founders for wheat.

### Comparison with Beagle

For the biparental cross scenarios without a pedigree, we compared the performance of our method (which we call AlphaPlantImpute2) with Beagle 5.1. Figure 5 shows the accuracy of AlphaPlantImpute2 versus the accuracy of Beagle 5.1. In all but one scenario, our method outperforms Beagle. As the number of individuals used to build the haplotype library increases, the accuracy of Beagle moves closer to that of our method, but only exceeds it for sugar beet with ten markers and 1000 individuals.

**Figure 5:**
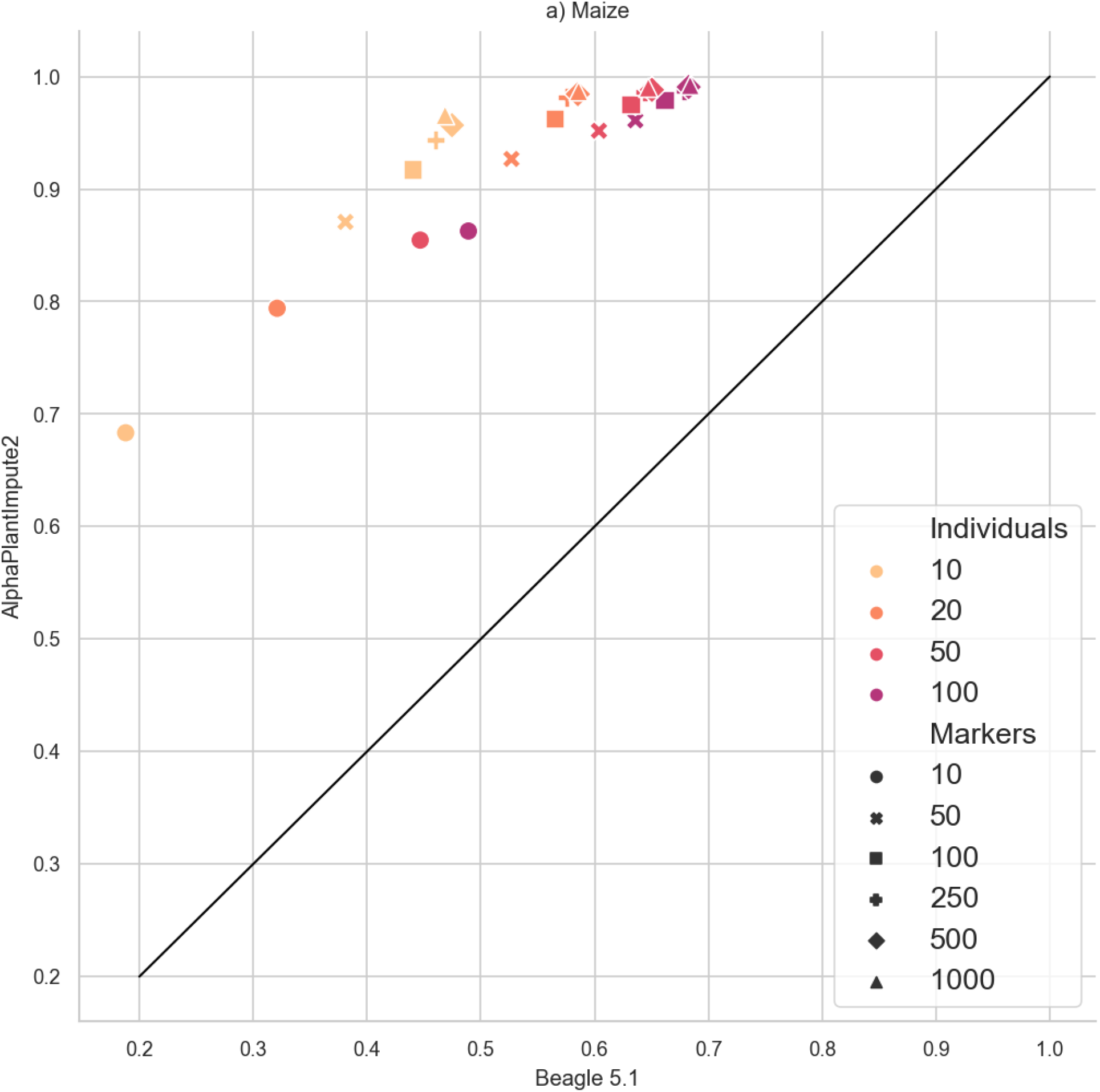
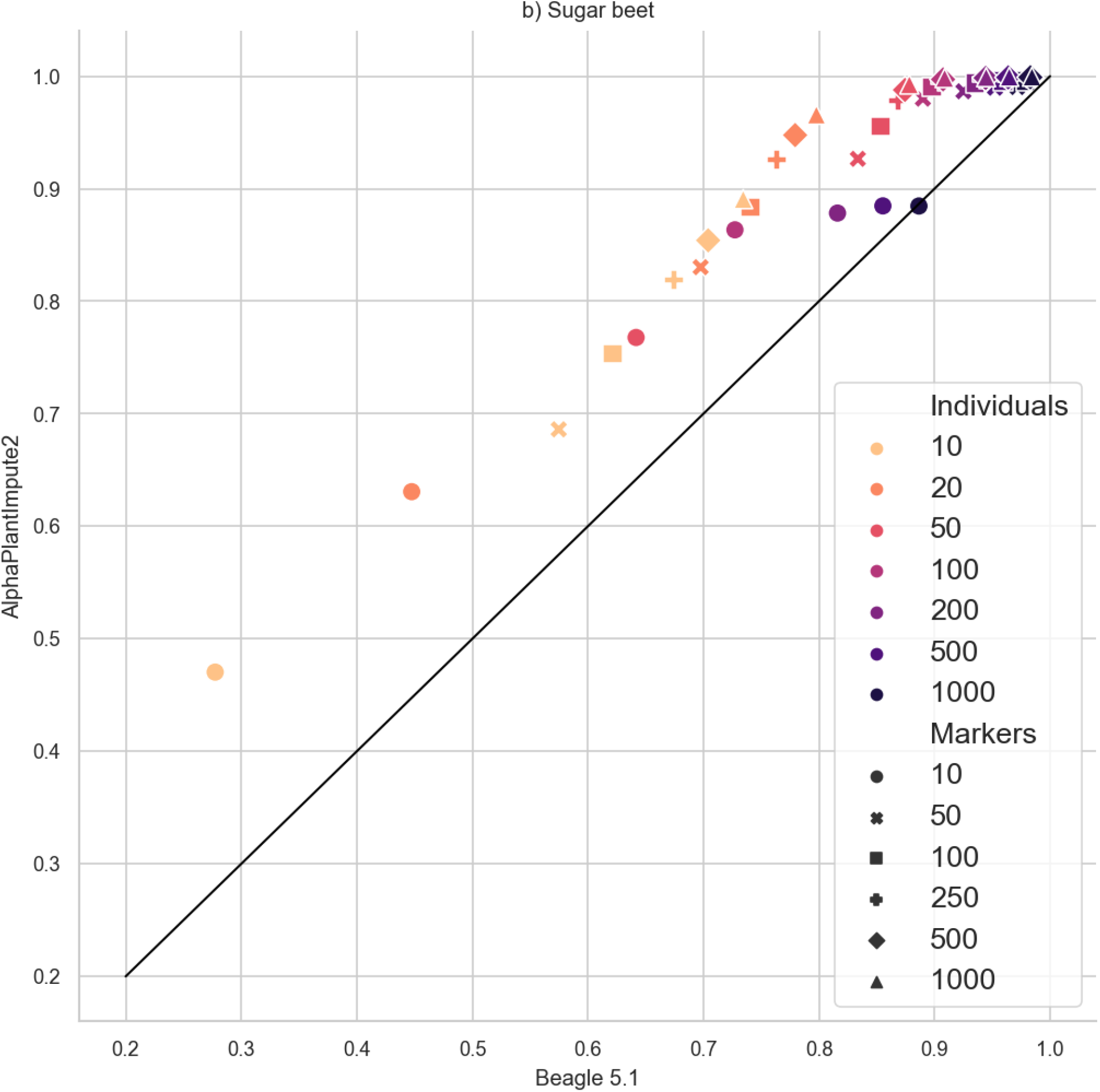
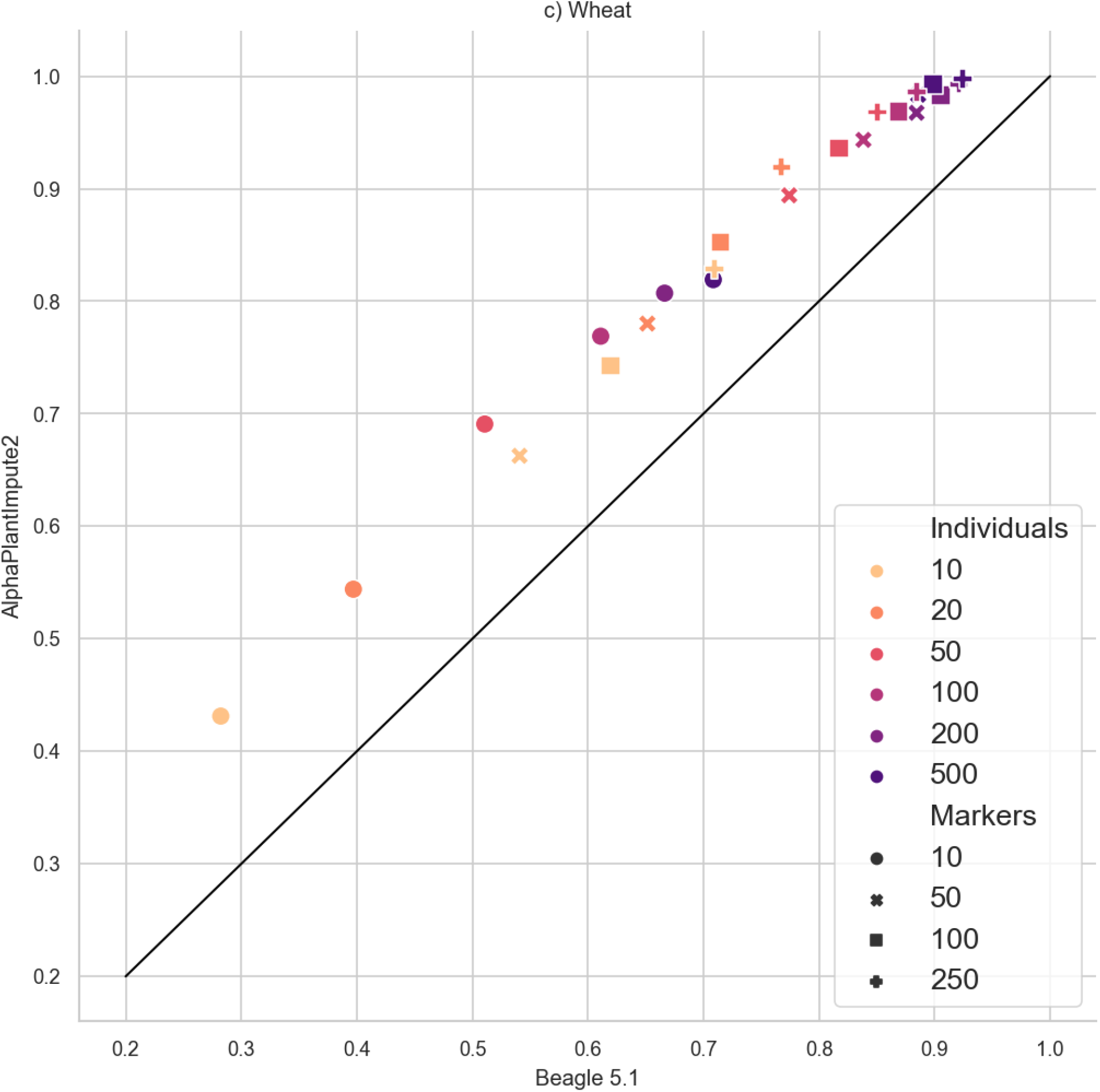
imputation accuracy of our method (AlphaPlantImpute2) versus Beagle 5.1 for biparental cross scenarios without a pedigree. Each point displays the median imputation accuracy over all imputed individuals, replicates and chromosomes. Model parameters, as described in the text.

## Discussion

In this paper, we demonstrate that plant breeders can effectively apply imputation in many crop species. Pedigree information is not required to obtain good accuracy, but it can improve the accuracy and the speed of imputation.

### Excellent imputation accuracy is possible without pedigrees

Our results show that excellent accuracy is possible in realistic scenarios. With 50 individuals genotyped at high-density and 250 markers per chromosome on the low-density array, imputation accuracies exceeded 0.97 for the three crop species. As the accuracies plateau around these values, there is an accuracy-cost trade-off that the plant breeder needs to consider. For example, doubling the number of individuals to 100 would increase the accuracy to > 0.99 for all crops, but doubles the genotyping cost.

### Pedigree information is useful

Pedigree information can increase accuracy by reducing the set of haplotypes that are used for imputation. Such a reduction also leads to a considerable speed improvement for the diploid hidden Markov model as the time it takes to impute individuals scales quadratically with number of haplotypes.

With relatively few founders, the imputation accuracy can be exceeded by scenarios without a pedigree when a greater number of progeny (compared to founders) genotyped at high-density are used to build the haplotype library. A possible strategy combining the best from scenarios with and without a pedigree would be to use founders *and* additional progeny genotyped at high-density to build the haplotype library. Then, as with the scenarios with a pedigree, only use the founder haplotypes to impute the progeny. Such a strategy would be an interesting follow on study.

## Conclusion

We have shown that accurate imputation is possible in commercial crop species. For the plant breeder, the choice of the number of markers on the low-density array, and the number of individuals genotyped at high-density is essential. Too few markers or individuals, and imputation accuracy will be low, but increasing the numbers provides ever-decreasing returns as the accuracies quickly plateau. For the crop species studied here, we found that excellent imputation accuracy needs approximately 50 individuals genotyped at high-density and low-density arrays of approximately 100 markers per chromosome. Accurate imputation does not require pedigree information or the genotyping of founders at high-density; however, these can easily be incorporated into the method and can increase imputation accuracy and computational speed.

## Acknowledgements

This work has made use of the resources provided by the Edinburgh Compute and Data Facility (ECDF) (http://www.ecdf.ed.ac.uk).

## Declarations

### Ethics approval

Not applicable

### Consent to participate

Not applicable

### Consent for publication

Not applicable

### Availability of data and material

Not applicable

### Availability of software

The software and source code is available from https://github.com/AlphaGenes/AlphaPlantImpute2.

### Author contributions

ST and AW designed and wrote the imputation software. ST, AW and JH designed the simulations. SK, MG, HSdS and VW provided the crop data and use cases for the software. ST ran the simulations and analysed the results. All authors contributed to writing the manuscript and approved the final manuscript

